# Hebbian induction adds AMPA-labile signaling units to CA3-CA1 synapses in the developing hippocampus

**DOI:** 10.1101/2024.09.25.614864

**Authors:** Therése Abrahamsson, Bengt Gustafsson, Eric Hanse

**Author notes:** Correspondence to: Eric Hanse Göteborg University, Department of Physiology Box 432, Medicinaregatan 11 405 30 Göteborg, Sweden.

## Abstract

In the 2^nd^ postnatal week hippocampus, Hebbian-induced long-term potentiation (LTP) of AMPA receptor-mediated transmission in CA3-CA1 synapses is not a genuine potentiation. Instead, it is a de-depression (unsilencing) and temporary stabilization of postsynaptically AMPA-labile synapses silenced by a prior test pulse (0.03 - 0.2 Hz) stimulation. In addition to such an LTP, Hebbian induction at these synapses also results in a labile potentiation that becomes depotentiated by test pulse stimulation, thus appearing as an Hebbian-induced short-term potentiation (STP). Although the induction of this labile potentiation was blocked in the combined presence of N-methyl-D-aspartate (NMDA) and metabotropic glutamate (mGlu) receptor antagonists, the depotentiation was not affected by these drugs. The labile potentiation was not associated with a change in paired-pulse ratio and was, after a depotentiation, fully re-established by a 20 min interruption of test pulse stimulation. These properties are shared with the silencing of previously non-stimulated (naïve) AMPA-labile synapses by such test pulse stimulation. However, the depotentiation following an Hebbian induction is not a re-silencing of naïve AMPA labile synapses since there is no correlation between the magnitudes of depotentiation and preceding silencing of naïve synapses. The present results suggest that Hebbian induction at these neonatal CA3-CA1 synapses, in addition to unsilencing and temporary stabilization of AMPA-labile transmission, creates a labile potentiation based on the insertion/activation of an additional AMPA-labile signaling unit to a pre-existing synapse.

## Introduction

In the neonatal brain Hebbian-induced synaptic plasticity is mainly an instrument for activity-dependent circuitry refinement (Katz & Shatz, 1996). When examined for neonatal hippocampal CA3-CA1 synapses, Hebbian-induced long-term potentiation (LTP) at these synapses also differs in several respects from the LTP found in the more adult synapses. Thus, while LTP at the more adult CA3-CA1 synapses relies on αCaMKII activation (Lisman *et al*., 2002; Nicoll & Schulman, 2023), LTP examined at postnatal days 7 - 9 (P7 - P9) does rather rely on activation of protein kinase A (PKA) (Yasuda *et al*., 2003). Moreover, while for the more adult LTP the AMPA receptor subunit GluA1 seems critically involved (Hayashi *et al*., 2000; Jensen *et al*., 2003; Chater & Goda, 2022), GluA2long and GluA4 subunits seem involved in early *N*-methyl-*D*-aspartate receptor (NMDAR) and PKA dependent synaptic AMPA receptor (AMPAR) incorporation (Zhu *et al*., 2000; Esteban *et al*., 2003; Kolleker *et al*., 2003; Qin *et al*., 2005; Luchkina *et al*., 2014). Furthermore, the neonatal LTP (<P12) is not a genuine potentiation but rather an AMPA unsilencing of AMPA labile synapses silenced by of a prior test pulse (0.03 - 0.2 Hz) stimulation (Abrahamsson *et al*., 2008; Cao & Harris, 2012). Hebbian induction thus stabilizes the AMPA receptor-mediated transmission at these synapses (Xiao *et al*., 2004; Abrahamsson *et al*., 2008), though not permanently (Strandberg & Gustafsson, 2024), indicating it rather selects which synapses that are to keep their AMPA signaling than to strengthen them.

Hebbian induction also produces a seemingly transient (5-20 min) potentiation that, in contrast to the LTP, transcends the naïve AMPA signaling level (Abrahamsson 2008), i.e., the level at which these synapses are all AMPA signaling (Xiao *et al*., 2004; Abrahamsson *et al*., 2008). Thus, this transient potentiation is a genuine potentiation of the AMPA receptor-mediated transmission. Moreover, it is not intrinsically transient but persists as long as the synapse is not activated (Strandberg & Gustafsson, 2024), i.e., it is transient because of the test pulse stimulation that follows the Hebbian induction event (Strandberg & Gustafsson, 2024). This Hebbian-induced potentiation thus seemingly responds to sparse synaptic activation in a similar manner as naïve (previously non-stimulated) neonatal CA3-CA1 synapses. The question then arises how the lability of this Hebbian-induced potentiation relates to the AMPA lability of the naïve synapses? In the present study, we have investigated properties of this Hebbian-induced potentiation of neonatal CA3-CA1 synapses from this perspective.

## Methods

### Slice preparation and solutions

Experiments were performed on hippocampal slices from 8 - 12 day–old Wistar rats. The animals were kept and killed in accordance with the guidelines of the Göteborg ethical committee for animal research. The rats were anaesthetized with isoflurane (Abbott) prior to decapitation. The brain was removed and placed in an ice–cold solution containing (in mM): 140 cholineCl, 2.5 KCl, 0.5 CaCl2, 7 MgCl2, 25 NaHCO3, 1.25 NaH2PO4, 1.3 ascorbic acid and 7 dextrose. Transverse hippocampal slices (300-400 µm thick) were cut with a vibratome (Slicer HR 2, Sigmann Elektronik, Germany) in the same ice–cold solution and they were subsequently stored in artificial cerebrospinal fluid (ACSF) containing (in mM): 124 NaCl, 3 KCl, 2 CaCl2, 4 MgCl2, 26 NaHCO3, 1.25 NaH2PO4, 0.5 ascorbic acid, 3 myo–inositol, 4 D,L– lactic acid, and 10 *D*–glucose. After 1 - 8 hours of storage at 25°C, a single slice was transferred to a recording chamber where it was kept submerged in a constant flow (∼2 ml minutes^-1^) at 30 – 32°C. The perfusion ACSF contained (in mM): 124 NaCl, 3 KCl, 4 CaCl2, 4 MgCl2, 26 NaHCO3, 1.25 NaH2PO4, and 10 *D*–glucose. Picrotoxin (100 µM) was always present in the perfusion ACSF to block GABAAR–mediated activity. All solutions were continuously bubbled with 95% O2 and 5% CO2 (pH ∼7.4). A cut between CA3 and CA1 and the higher than normal Ca^2+^ and Mg^2+^ concentrations were used to prevent spontaneous network activity. Under these conditions the spontaneous synaptic activity in the slice preparation is very low, spontaneous excitatory postsynaptic currents (EPSCs) occurring at a frequency of about 0.3 – 1 Hz (Hsia *et al*., 1998; Groc *et al*., 2002).

### Recording and analysis

Electrical stimulation of Schaffer collateral afferents was carried out in the stratum radiatum. Stimuli consisted of biphasic constant current pulses (200 + 200 µs, STG 1004, Multi Channel Systems MCS Gmbh, Reutlingen, Germany) delivered through either a glass pipette (resistance ∼ 0.5 - 1 MΩ) or an insulated tungsten microelectrode (resistance ∼ 0.3 - 0.5 MΩ). Field EPSP (fEPSP) recordings were made by means of a glass micropipette (∼ 1 MΩ, filled with 1 M NaCl) in the stratum radiatum. Perforated patch–clamp recordings were performed on visually identified pyramidal cells, using infrared–differential interference contrast videomicroscopy mounted on a Nikon E600FN microscope (Nikon, Japan). The pipette solution contained (in mM): 130 KCl, 2 NaCl, 20 HEPES, 0.2 EGTA, 4 Mg-ATP, 0.4 GTP and amphotericin B (240 µg/ml) (pH ∼7.3 and osmolality 270–300 mOsm). Lucifer yellow (0.05%) was always present in the perforating solution to detect possible membrane leakage using a fluorescence microscope. If lucifer yellow entered the cell the experiment was discarded. EPSCs and fEPSPs were recorded at a sampling frequency of 10 kHz and filtered at 1 kHz, using an EPC–9 amplifier (HEKA Elektronik, Lambrecht, Germany). For AMPA EPSC recordings cells were held in voltage-clamp mode at –70 mV. Series resistance was monitored using a 5 ms 10 mV hyperpolarizing pulse and it was not allowed to change more than 20%, otherwise the recording was not included in the analysis.

fEPSPs and EPSCs were analyzed off–line using custom–made IGOR Pro (WaveMetrics, Lake Oswego, OR) software. fEPSPs and EPSCs amplitudes were measured as the difference between the baseline level immediately preceding the stimulation artefact, and the mean amplitude during a 2 ms time window around the negative peak between 3 and 8 ms after the stimulation artefact. fEPSPs were also estimated by linear regression of the initial slope. Since our experimental design precludes adjustment of stimulation intensity (Abrahamsson *et al*., 2007) experiments in which the naïve fEPSPs exhibited signs of population spike activity were discarded. For the fEPSPs not excluded by this criterion initial slope and amplitude measurements gave the same results. Since the amplitude measurements were less noisy these measurements were used for illustrations. The presynaptic volley was measured as the amplitude of the initial positive–negative deflection and was not allowed to change more than 10%, otherwise the experiment was discarded. Data are expressed as means ± SEM.

### Drugs

Chemicals were from Sigma–Aldrich (Stockholm, Sweden) except for *D*-AP5 and LY 341495 (Tocris Cookson, Bristol, UK).

## Results

### Hebbian-induced transient potentiation at neonatal CA3-CA1 synapses

When previously non-stimulated (naïve) CA3-CA1 synapses in slices from P8-P12 rats are tetanized, there is no succeeding LTP but only a transient potentiation decaying within 5-20 min when subjected to test pulse stimulation (Abrahamsson *et al*., 2008). This is illustrated in Fig. 1A where a high frequency stimulation (HFS; three 20-impulse, 50 Hz, trains at 0.05 Hz) used as an Hebbian induction event is given after that the naïve fEPSP level was established by using two test pulse stimulations prior to the tetanization. When instead allowing for an initial period of test pulse stimulation (0.2 Hz), the fEPSP becomes depressed and a subsequent HFS results in an apparent LTP preceded by a transient potentiation decaying to the naïve fEPSP level (Fig. 1B). Under these conditions the transient potentiation reached a peak value of 131 ± 3.3% (n = 23), of the naïve fEPSP level which is comparable to the peak potentiation of 136 ± 3.1% (n = 6) of the naïve level shown in Fig. 1A. When plotting superimposed, the test pulse-induced depression from the naïve level (Fig. 1A) and that from the peak of the potentiation (Fig. 1B), these time courses well overlap (Fig. 1C).

**Figure 1.**
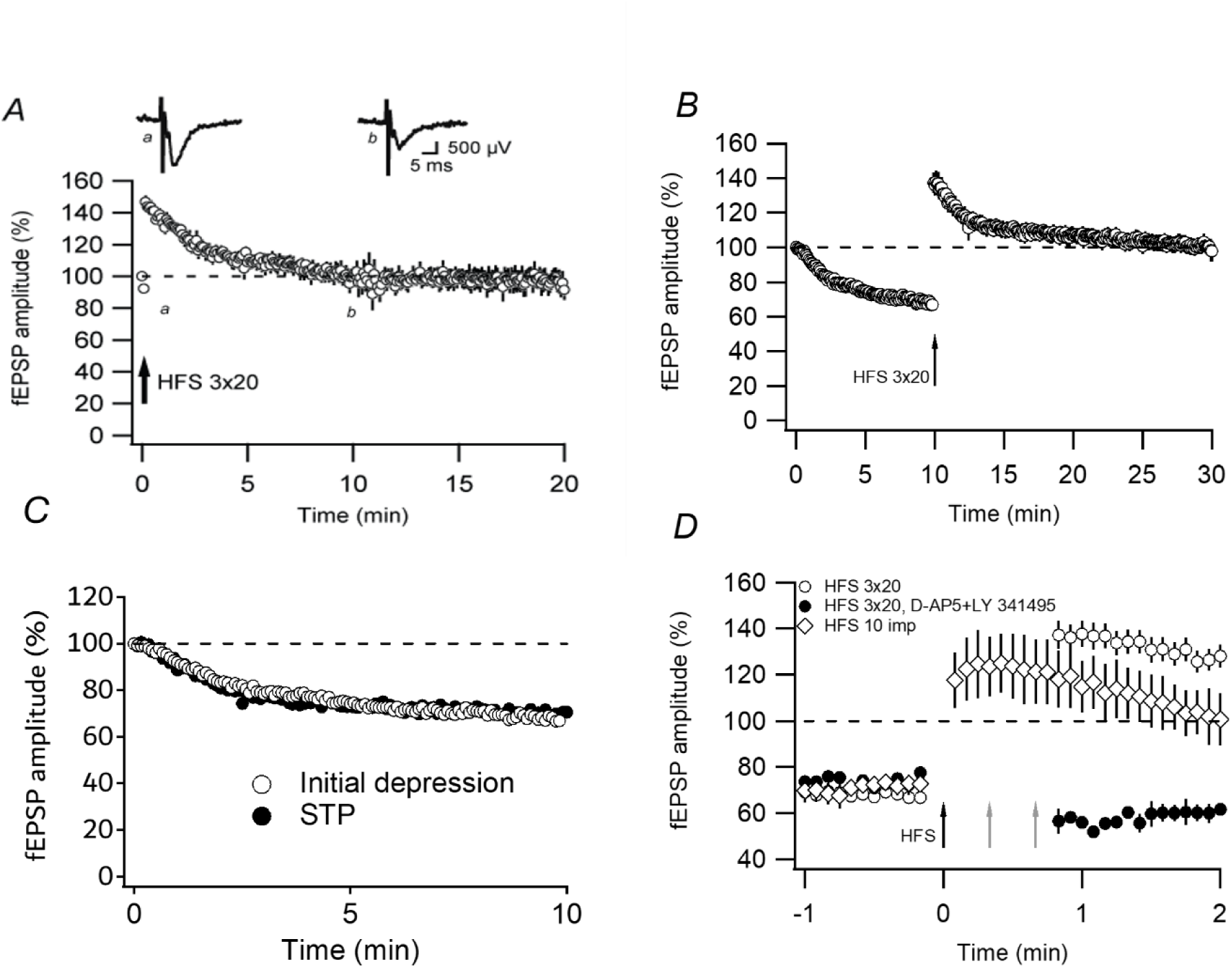
Hebbian-induced transient potentiation. A, after establishing the naïve level by two test stimuli, the synapses were subjected to high-frequency stimulation (HFS) (20-impulse trains at 50 Hz, repeated three times 20 s apart (n = 6 experiments)). The potentiation is expressed as percent of the naïve level. Test stimulus frequency was 0.2 Hz. B, the same as in A but the HFS was preceded by 10 min of test pulse stimulation. Example field EPSPs (average of 10 records) in A and B are taken at time points indicated in the figure. C, onset of the transient potentiation. The HFS-induced potentiation shown in B is re-plotted on an expanded time scale (open circles) together with the HFS-induced depression observed in the presence of the NMDA receptor antagonist D-AP5 (50 µM) and mGluR antagonist LY 341495 (50 µM) (closed circles, n = 5). Included is also the potentiation produced by a single 10-impulse tetanus (50 Hz) (diamonds, n = 5).

When performing the experiment in Fig. 1B in the presence of NMDA and mGlu receptor antagonists the HFS produces essentially no transient component when measured from 10 s after the (third) tetanization event (Fig. 1D, closed circles). This result suggests that none of the observed potentiation obtained in the absence of these antagonists (Fig. 1B and D, open circles) reflects a presynaptically based plasticity, such as augmentation or post-tetanic potentiation. Following a brief single tetanus (10-impulse, 50 Hz) to these neonatal CA3-CA1 synapses, a significant potentiation is seen already within the first 10 seconds post-tetanus, and a maximal potentiation 30 seconds post-tetanus (Fig. 1D, open rhombi). The Hebbian-induced potentiation of these neonatal synapses is thus established within half a minute, which is similar to what has been shown for the Hebbian-induced NMDA receptor dependent potentiation of more adult hippocampal synapses (Gustafsson *et al*., 1989; Hanse & Gustafsson, 1992). The 2^nd^ and the 3^rd^ tetanization (see arrows in Fig. 1D) thus occur at fully potentiated synapses without this condition apparently contributing to any stabilization of the potentiation.

### The transient potentiation is a labile potentiation that decays in a stimulation-dependent, but NMDAR/mGluR-independent, manner

When the test pulse stimulation was interrupted for five minutes immediately after the HFS, the peak potentiation was 52 ± 6.3% of the naïve level (n = 4) when stimulation was resumed (Fig. 2A), a potentiation more than comparable with the peak potentiations observed in the absence of such stimulus interruption (Fig. 1). This result agrees with a previous study of these synapses that the potentiation persists during stimulus interruption (for at least a few hours) {Strandberg, 2024 #15}, and becomes depotentiated by the test pulse stimulation. The above experiment was also performed applying NMDA and mGlu receptor antagonists immediately after the HFS (Fig. 2B), demonstrating a similar retainment of potentiation by stimulus interruption and a subsequent stimulation dependent depotentiation. Thus, when measured one and ten minutes after the resumption of stimulation, the fEPSP was 144 ± 12% and 112 ± 12% (n = 3), respectively, of the naïve level (Fig. 3B), compared with 152 ± 6.3% and 112 ± 2.5%, respectively, in the absence of these antagonists (Fig. 2A). The stimulation dependent, rather than time dependent, decrease of the labile potentiation was also apparent when reducing the test pulse frequency from 0.2 Hz to 0.05 Hz (Fig. 2C). In fact, in agreement with test pulse-induced depression from the naïve level {Strandberg, 2009 #52;Xiao, 2004 #14}, the 0.2 Hz and 0.05 Hz-induced depotentiations are well superimposed when plotted against number of stimuli (Fig. 2D) rather than time (Fig. 2C),

**Figure 2.**
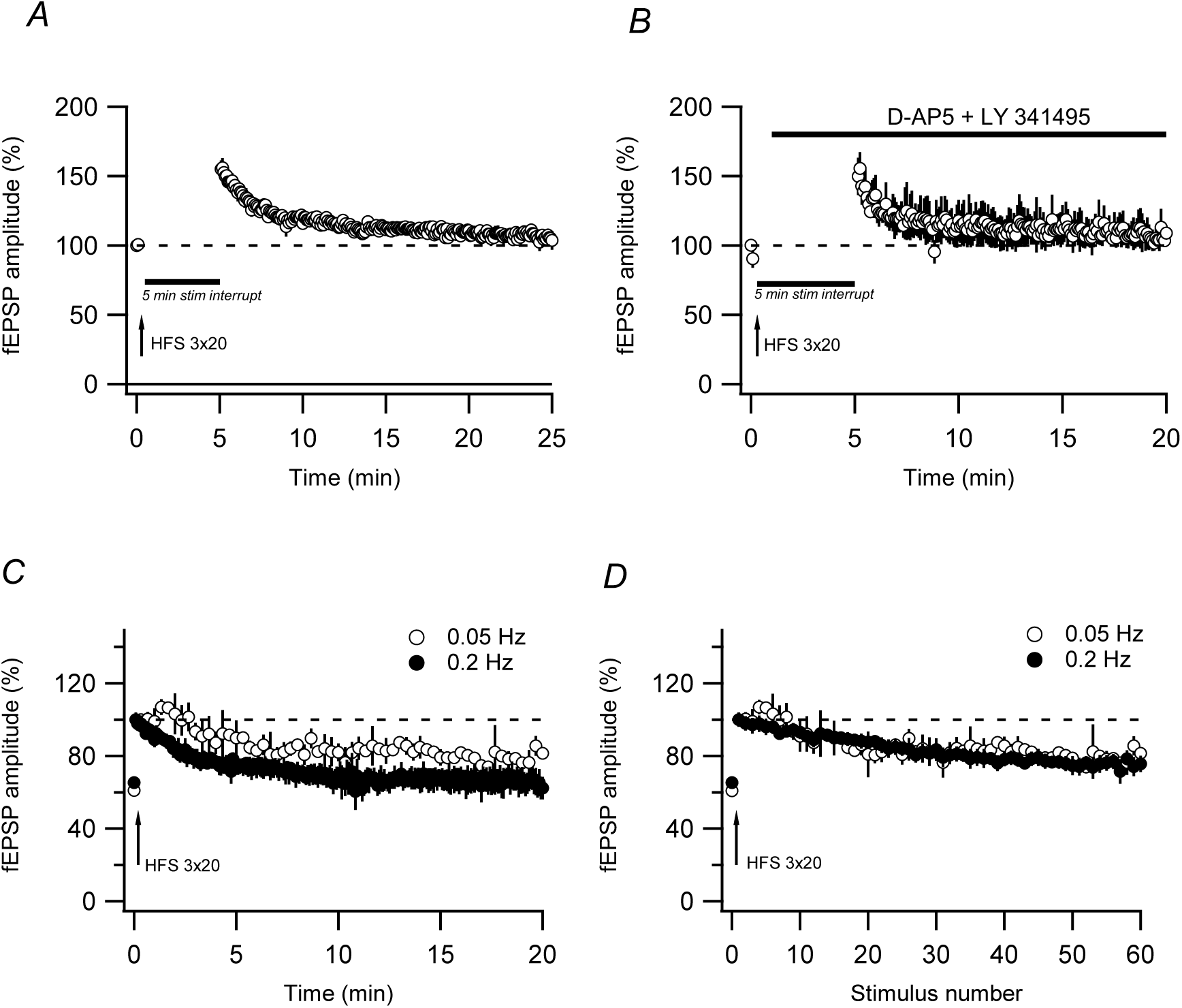
The transient potentiation decays in a stimulation-dependent, but NMDAR/mGluR- independent, manner. A, test pulse stimulation (0.2 Hz) was interrupted immediately following an HFS-induced potentiation (n = 4). and re-started 5 min later. B, same as in A, but combined with an application of D-AP5 (50 µM) and LY 341495 (50 µM) immediately following the tetanization (n = 3).

**Figure 3.**
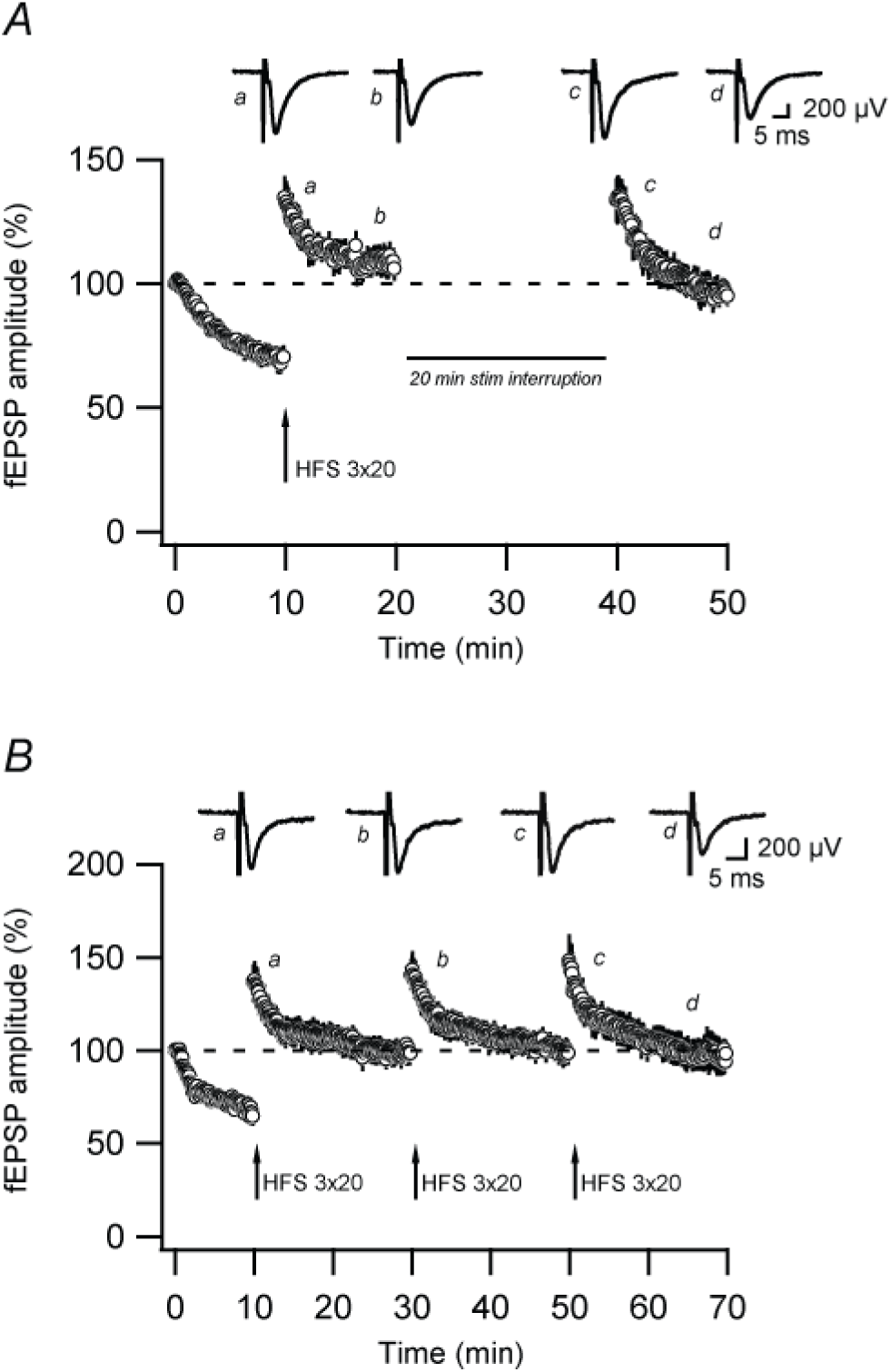
The labile potentiation is restored by stimulus interruption and by repeated tetanization. A, an HFS-induced potentiation is preceded by 10 min of test pulse (0.2 Hz) stimulation (n = 4). Ten min after the HFS, the test pulse stimulation was interrupted for 20 min, allowing for recovery of the labile potentiation. B, an HFS, as in A, was repeated three times with 20 min intervals (n = 5).

### The labile potentiation is restored by stimulus interruption and by renewed tetanization

Test pulse-induced depression of naïve synapses following a limited number of stimulations (≤100) is not stable, but when the test pulse stimulation is interrupted the fEPSP recovers to the naïve level within ∼20 minutes (Abrahamsson *et al*., 2007). That is, synapses silenced in this manner must be activated for the silent state to remain. To test whether the labile potentiation also exhibits this feature, test pulse stimulation was interrupted for 20 minutes after that the potentiation had decayed to the naïve level. This stimulus interruption resulted in a complete re-establishment of the potentiation (Fig. 3A). The peak of the control and of the re-establishment potentiations were 132 ± 8.2% and 134 ± 8.7% (n = 4), respectively, of the naïve level. In separate experiments, a briefer (5 min) stimulus interruption resulted in a partial re-establishment, the peak potentiation being 123 ± 5.7% (n = 3) of the naïve level (not shown). This partial re-establishment, following 5 min of stimulus interruption, then indicates a similar time course of recovery by stimulus interruption as that of test pulse-induced depression of naïve synapses (Abrahamsson *et al*., 2007). As shown previously (Abrahamsson *et al*., 2008), when instead repeating the HFS following 20 min of test pulse stimulation the subsequent potentiation did not differ from that obtained after the first HFS (Fig. 3B). Thus, the labile potentiation induced by an HFS can equally well be restored either by 20 min of stimulus interruption or by a renewed HFS.

### The labile potentiation is not a repetition of the test pulse-induced depression of naïve synapses

The similarity between the test pulse-induced depression from a naïve level and that from the post-tetanus level (Fig. 1C), raises the question whether these depressions are actually the same. That is, the HFS induces a separate lasting potentiation process, and the stimulation dependent depression after the HFS is actually that of naïve synapses. If so, examining the individual experiments, such as those in Fig. 1B, one might expect correlations between the magnitude of depression observed prior to the HFS and that observed after that event. However, when the peak magnitude of the potentiation was compared with that of the preceding test pulse-induced depression, there was no significant correlation between these magnitudes (r = 0.08, p > 0.05, n = 23; Fig. 4A). Moreover, when measured 20 min after the HFS, there was not either any significant correlation between the deviation from the naïve level and the amount of preceding test pulse-induced depression (r = 0.34, p > 0.05, n = 23; Fig. 4B). When comparing experiments selected for the largest and smallest magnitudes of preceding test pulse-induced depression (35.6 ± 0.8%, n = 5 (Fig. 4C) and 23.0 ± 1.6%, n = 5 (Fig. 4D), respectively), the magnitudes of the potentiation did not significantly differ (35.7 ± 7.3% and 41.2 ± 8.9%, respectively, p > 0.05). Similarly, when selecting experiments with the largest and the smallest magnitudes of the potentiation (41.7 ± 7.7%, n = 5 (Fig. 4E) and 19.6 ± 5.2%, n = 5 (Fig. 4F), respectively, p < 0.05), the amount of preceding test pulse-induced depression (68.8 ± 4.3% and 67.9 ± 2.5%, respectively, p > 0.05) did not differ. It would thus appear that the labile potentiation observed after the HFS reflects a process not present prior to this induction.

**Figure 4.**
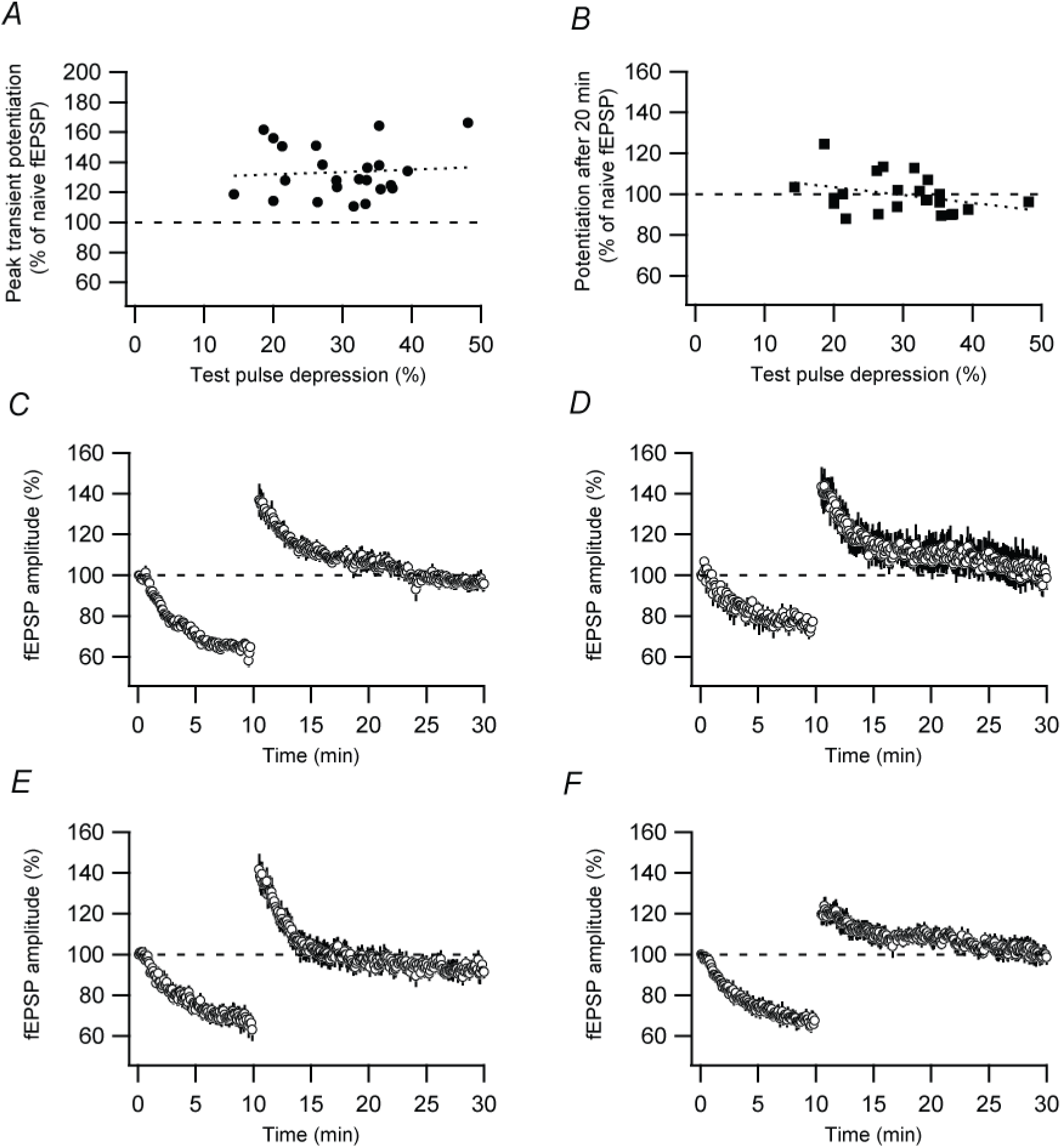
Lack of correlation between the labile potentiation and the test pulse-induced depression of naïve synapses. A, the peak value of the HFS-induced potentiation exceeding the naïve level is plotted against the amount of preceding test pulse-induced depression. B, the difference between the HFS-induced potentiation (measured at 20 min after tetanization) and the naïve level, is plotted against the amount of preceding test pulse-induced depression. C-F, following 10 min of test pulse stimulation (0.2 Hz) an HFS was given. Experiments included in C (n = 5) and D (n = 5) are those preceded by the largest and smallest test pulse-induced depressions, respectively. Experiments included in E (n = 5) and F (n = 5) are those with the largest and smallest transient potentiations, respectively.

### Neonatal labile potentiation is not associated with change in paired-pulse plasticity

To address whether the labile potentiation reflects a presynaptic change in release probability, we examined for possible changes in the paired-pulse ratio during the potentiation. For these experiments intracellular recording rather than field recording was used to avoid the induction of a paired-pulse stimulation-induced NMDAR/voltage-gated calcium channel dependent depression in these developing synapses (Wasling *et al*., 2002). Since the induction of Hebbian plasticity may be compromised in the whole-cell recording mode, the perforated patch-clamp mode was used. HFS (paired with a depolarizing pulse to 0 mV for 400 ms) was applied after 5- 12 minutes of test stimulation consisting of paired-pulse stimulation (50 ms interval) at 0.2 Hz. In conformity with results obtained for test pulse-induced depression of naïve synapses (Xiao *et al*., 2004; Abrahamsson *et al*., 2005), the potentiation was not associated with any significant change in the paired-pulse ratio (Fig. 5A, B).

**Figure 5.**
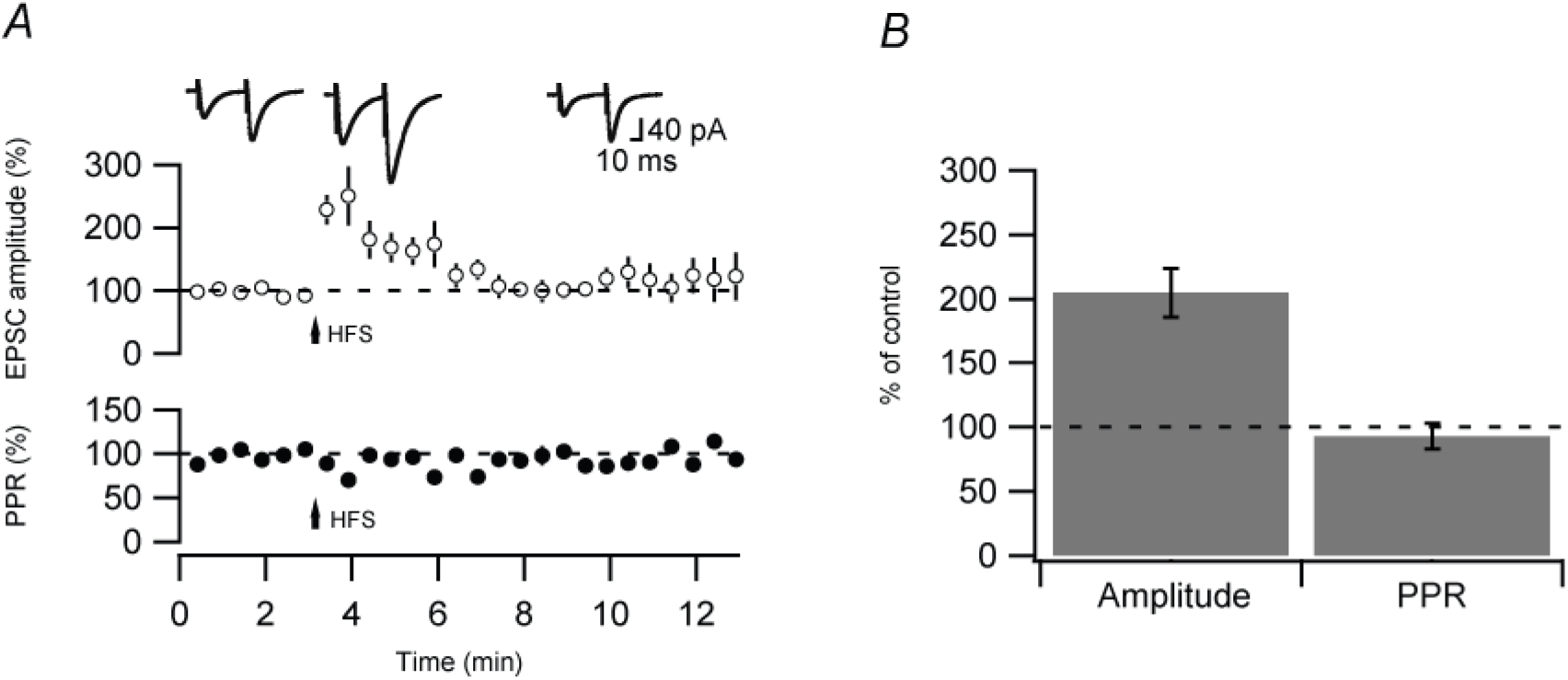
Labile potentiation is associated with unchanged paired-pulse ratio. A, a 20-impulse train (50 Hz) was paired with a 400 ms depolarizing pulse to 0 mV, repeated three times 20 s apart, using perforated patch-clamp recordings (n = 7). As noted in the graph, this induction event resulted largely in a transient potentiation (open circles) and no associated change in the paired-pulse ratio (closed circles). Test pulses consisted of paired-pulse stimuli (50 ms interval) at 0.2 Hz, this stimulation starting 2-9 minutes prior to the Hebbian induction event (see arrow), i.e., the synapses were at time zero in the graph not naïve. Each point in the graph is the average of six successive stimuli. B, Bar diagram showing a comparison between the peak of the transient potentiation (∼1 minute after the induction event) and the corresponding change in paired-pulse ratio. Example EPSCs (average of 6 records) in A are taken at time points indicated in the figure.

## Discussion

In agreement with a recent study of 2^nd^ postnatal week hippocampal CA3-CA1 synapses (Strandberg & Gustafsson, 2024) we find that the Hebbian-induced transient potentiation is not inherently transient but is a labile potentiation depotentiated by even sparse synaptic activity. In fact, in the absence of such activity this potentiation has been found to be stable for hours (Strandberg & Gustafsson, 2024). The present results further reinforce the similarity between the lability of this potentiation and the lability of previously non-stimulated naïve synapses. Thus, the depotentiation follows the same stimulation dependent course as the depression of naïve synapses, it does not require NMDAR or mGluR activation, it is not associated with a change in the paired-pulse ratio, and it is, following a limited number of stimulations, fully reversed by 20 min of stimulus interruption (Xiao *et al*., 2004; Abrahamsson *et al*., 2007; Strandberg & Gustafsson, 2024).

A simple explanation for these common properties is that this labile potentiation actually reflects the test pulse-induced depression observed prior to the HFS. That is, the HFS produces a genuine stable LTP process unrelated to a prior test pulse-induced depression, as well as de- depresses the test pulse depressed synapses. These latter synapses then subsequently again become depressed by the post-HFS test pulse stimulation. However, we found among the experiments no correlation between the peak magnitude of the potentiation and the magnitude of the preceding test pulse-induced depression. We also found the naïve level to be an approximate ceiling for an apparent LTP, independent of the magnitude of the preceding test pulse-induced depression, as expected if this apparent LTP is a de-depression of the prior test pulse-induced depression. None of these results supports the notion that the labile potentiation should reflect the lability of naïve synapses but rather favor the notion that the labile potentiation is explained by the postsynaptic addition of AMPA labile transmission not present prior to the Hebbian induction event {Xiao, 2004 #14}.

It should be noted that potentiations that are preserved in the absence of activity, and eliminated by sparse activity, have been previously observed in the hippocampus. The mossy fiber-CA3 synapses exhibit a NMDAR independent post-tetanic potentiation whose lifetime is extended in the absence of presynaptic activity (Vandael *et al*., 2020). A post-tetanic potentiation with similar properties has also been described among young adult CA3-CA1 synapses (Pradier *et al*., 2018). In contrast to the presently observed potentiation this post-tetanic potentiation did not became re-established by stimulus interruption. In addition, Volianskis and Jensen (Volianskis & Jensen, 2003) has shown that NMDAR-dependent short-term potentiation at adult CA3-CA1 synapses is preserved in the absence of presynaptic activity. Thus, NMDAR-dependent short-term potentiation in neonatal CA3-CA1 synapses (present results) share this property with its more adult counterpart. However, it is not known whether the adult short-term potentiation become re-estblished by stimulus interruption. Also, in contrast to the present results, the adult short-term potentiation was found to be associated with paired-pulse changes (Volianskis & Jensen, 2003).

### Postsynaptic expression for the labile potentiation

Much evidence has been presented linking the test pulse-induced depression of naïve synapses to a postsynaptic AMPA silencing (Xiao *et al*., 2004; Abrahamsson *et al*., 2005; Wasling *et al*., 2012). The general similarity between the depotentiation of the labile potentiation and the test pulse-induced depression of naïve synapses makes it reasonable to propose that the labile potentiation is postsynaptically expressed, and explained by the insertion and subsequent removal of AMPARs. We found that the peak magnitude of the labile potentiation did not match quantitatively the amount of preceding depression. This result suggests that the labile potentiation is specific neither for the synapses that were silenced by the preceding test pulse stimulation nor for the synapses that were not silenced by that stimulation. The labile potentiation may then be explained by the new insertion of AMPARs into any, or all, of the synapses exposed to the HFS, expanding the number of AMPARs at the postsynaptic location.

In the 2^nd^ postnatal week the CA3-CA1 synapse only contains a single release site (Hsia *et al*., 1998; Groc *et al*., 2002), likely representing a single nanocolumn (Sakamoto *et al*., 2018; Ramsey *et al*., 2021; Hruska *et al*., 2022), an integrated subsynaptic component for vesicle release and transmitter reception. The test pulse-induced silencing of naïve synapses is thus the full removal of the AMPARs from the single nanocolumn of some of these synapses, creating AMPA-silent synapses. Considering the relative ineffectiveness of adding AMPARs to a given nanocolumn (Sinnen *et al*., 2017), we speculate that the Hebbian-induced addition of AMPARs is that of an added nanocolumn, these AMPARs being labile. The depotentiation would then mechanistically be similar to the depression of a naïve synaptic input, i.e., the stimulation dependent full removal of AMPARs from a nanocolumn.

Since an Hebbian induction stabilizes naïve AMPA labile synapses (Xiao *et al*., 2004; Abrahamsson *et al*., 2008), although not permanently (Strandberg & Gustafsson, 2024), there has to be a distinction between the original and the added nanocolumns, either with respect to the AMPARs themselves or to AMPAR-associated proteins, keeping the added AMPARs labile. In the 2^nd^ postnatal week the AMPARs in naïve synapses are likely of the GluA4/ GluA2long subtypes, receptors that can be inserted via PKA activation alone. The neonatal LTP, as well as the labile potentiation, is also mimicked by the application of the PKA activator forskolin (Yasuda *et al*., 2003; Abrahamsson *et al*., 2008). The insertion of such AMPARs would then imply that the lability does not reside in the inserted AMPARs themselves, but in differential properties of AMPAR-associated proteins.

A remarkable feature of the labile potentiation is that it became re-established fully by 20 min of stimulus interruption. That is, while this potentiation reflects a HFS-induced potentiation process that is maintained at least for hours unless the synapse is stimulated, it can also after its depotentiation become repotentiated by stimulus interruption alone. In this respect, the depotentiated synapse resembles the silenced synapse that also can regain AMPA signaling by stimulus interruption alone (Abrahamsson *et al*., 2007). As noted above, AMPARs containing GluA4 or GluA2long subunits can be synaptically inserted by spontaneous activity itself in an NMDAR- and PKA-dependent manner (Zhu *et al*., 2000; Kolleker *et al*., 2003). However, while the silenced synapse returns to a naïve state existing prior to the test pulse stimulation, the repotentiation of the depotentiated synapse is to a state that had required an HFS for its induction. However, if, as speculated above, the labile potentiation reflects the insertion of an additional nanocolumn with labile postsynaptic AMPA signaling, the distinction may be more apparent than real as long as the added nanocolumn remains.

The restoration of AMPA signaling in existing, or newly established, nanocolumns by inactivity may signify a homeostatic plasticity. It is then noteworthy that our present results indicate that the very same synaptic modification can be induced either by Hebbian activity, or as a consequence of a homeostatic response to inactivity. A similar convergence between the expression of homeostatic and Hebbian plasticity appears also to exist in more mature CA3-CA1 synapses where added nanocolumns to existing synapses have been implicated for homeostatic plasticity induced by partial AMPAR blockade (Chipman *et al*., 2022) and for NMDAR-dependent long-term potentiation (Sinnen *et al*., 2017; Choquet & Hosy, 2020; Hruska *et al*., 2022). The time requirement for induction differs however, such that the Hebbian plasticity develops within tens of seconds (Gustafsson *et al*., 1989) (present results) whereas the homeostatic counterpart develops within minutes of inactivity (Chipman *et al*., 2022) (present results).

### Functional considerations

The lability of this potentiation has also implications under which functional conditions it will be observed. Thus, when LTP is induced during low frequency activity of a synapse paired with strong depolarization of the postsynaptic neuron, one might expect this labile potentiation to quickly vanish. In fact, the initial transient phase of potentiation characteristically observed when LTP is induced by a brief tetanization is largely absent following LTP induction using low-frequency pairing (Isaac *et al*., 1995; Liao *et al*., 1995; Durand *et al*., 1996; Rumpel *et al*., 1998; Montgomery *et al*., 2001). Likewise, the transient potentiation observed following forskolin application was not observed unless stimulation was interrupted during the wash-in of the drug (Abrahamsson *et al*., 2008).

As suggested above, the labile potentiation is not mechanistically distinct from developmental LTP but rather mechanistically related, being explained by the insertion of an additional nanocolumn. Considering that adult LTP may be explained by an increased number of nanocolumns in a synapse (Sinnen *et al*., 2017; Choquet & Hosy, 2020; Hruska *et al*., 2022), the labile potentiation in these neonatal synapses may be an unstable form of the adult LTP which starts to appear after P12 (Abrahamsson *et al*., 2008). The labile behavior may then be related to the absence, prior to P12, of e.g. αCaMKII activity and/or of scaffolding proteins such as PSD-95/PSD-93 that may be necessary for insertion of new stable AMPA clusters into the nanocolumns. The labile potentiation observed in these neonatal synapses may then be a forerunner to the LTP to be when these synapses operate in a more mature hippocampus.

